# Identification of genes required for apical protein trafficking in Drosophila photoreceptor cells

**DOI:** 10.1101/653469

**Authors:** Azadeh Laffafian, Ulrich Tepass

**Author notes:** **Address for correspondence:** Ulrich Tepass, Department of Cell and Systems Biology University of Toronto, 25 Harbord Street, Ontario M5S 3G5, Canada.

## Abstract

Drosophila photoreceptor cells (PRCs) are highly polarized epithelial cells. Their apical membrane is further subdivided into the stalk membrane and the light-sensing rhabdomere. The photo-pigment Rhodopsin1 (Rh1) localizes to the rhabdomere, whereas the apical determinant Crumbs (Crb) is enriched at the stalk membrane. The proteoglycan Eyes shut (Eys) is secreted through the apical membrane into an inter-rhabdomeral space. Rh1, Crb, and Eys are essential for PRC development, normal vision, and PRC survival. Human orthologs of all three proteins have been linked to retinal degenerative diseases. Here, we describe an RNAi-based screen examining the importance of approximately 240 trafficking-related genes in apical trafficking of Eys, Rh1, and Crb. We found 28 genes that have an effect on the localization and/or levels of these apical proteins and analyzed several factors in more detail. We show that the Arf GEF protein Sec71 is required for biosynthetic traffic of both apical and basolateral proteins, that the exocyst complex and the microtubule-based motor proteins dynein and kinesin promote the secretion of Eys and Rh1, and that Syntaxin 7/Avalanche controls the endocytosis of Rh1, Eys, and Crb.

**Article summery:** Photoreceptor cells (PRCs) rely on polarized vesicle trafficking to deliver key secreted and transmembrane proteins to their correct locations. Failure to do so causes defects in PRC development, function, and survival leading to retinal disease. Using the fruit fly Drosophila as a model we have identified 28 genes that are required for the trafficking of the three apical proteins Rhodopsin 1, Crumbs, and Eyes Shut. Human homologs of all three genes are associated with retinal degeneration. We characterized several genes to reveal novel mechanisms of vesicle trafficking in photoreceptor cells at different points in the biosynthetic or endocytotic pathways.

## Introduction

Drosophila photoreceptor cells (PRCs) are an important model for the epithelial differentiation of a sensory cell and to study vesicle trafficking and neuro-degeneration (for reviews see Tepass and Harris, 2007; Shieh, 2011; Xiong and Bellen, 2013; Schopf and Huber, 2017). PRCs have specialized apical and basolateral membranes that are segregated by an epithelial adherens junction, a zonula adherens. While the basolateral membrane extends an axon, the apical membrane differentiates a light sensing organelle, the rhabdomere. In addition to the rhabdomere, the apical membrane of PRCs contains the stalk membrane domain that connects the rhabdomere to the zonula adherens. Here, we have identified factors that contribute to the trafficking of three proteins - Rhodopsin 1 (Rh1), Crumbs (Crb), and Eyes shut (Eys) to the apical membrane of PRCs to further our understanding how the vesicle trafficking machinery contributes to the maintenance and function of a complex epithelial sensory cell.

Rhodopsin photopigments are seven-pass trans-membrane G-protein-coupled receptors localized at rhabdomeres, an array of 10th of thousands tightly packed microvilli. The main rhodopsin protein, Rh1, is found in PRCs R1 to R6. Mutations in rhodopsin and defects in its trafficking are the most frequent cause of retinal degenerative diseases in flies and humans (Colley et al., 1995; Xiong and Bellen, 2013; Nemet et al., 2015).

The apical polarity determinant Crumbs (Crb) is found at the stalk membrane. This transmembrane protein is important in regulating the length of the stalk; a reduction of Crb limits and an overexpression of Crb expands the stalk membrane (Pellikka et al., 2002). Crb is also important in the maintenance of the zonula adherens, and the proper distal to proximal elongation of rhabdomeres (Pellikka et al., 2002; Izaddoost et al., 2002). *crb* mutant PRCs show light-induced degeneration (Johnson et al., 2002) and a human homologue of Crb (CRB1) has been linked to the retinal degenerative diseases, retinitis pigmentosa (RP12) and Leber congenital amaurosis (LCA) (den Hollander et al., 1999, Richard et al., 2006; den Hollander et al., 2008; Bujakowska et al., 2012; Pellikka and Tepass, 2017).

The apical membranes of PRCs face a luminal space called the interrhabdomeral space (IRS). The IRS is important in vision in flies as it physically separates the rhabdomeres within one ommatidium from each other. The proteoglycan Eys shut (Eys) is essential in the formation of the IRS (Husain et al., 2006; Zelhof et al., 2006). Eys is thought to be secreted through the stalk membrane into the IRS (Husain et al., 2006). Similar to mutations in rhodopsin and CRB1, also mutations in the human EYS have been linked to retinal degenerative diseases such as retinitis pigmentosa (Abd El-Aziz et al., 2008; Collin et al., 2008).

The fact that several apical transmembrane or secreted proteins play key roles in PRC development and disease motivates a careful assessment of the mechanism that transport and target these proteins. Several factors have been identified that are involved in apical trafficking in Drosophila PRCs (see Figure 1). Examples include Rab1 and Syntaxin 5 (Syx5) that are essential in ER to Golgi trafficking (Satoh et al., 1997; Satoh et al., 2016). Rab6 is important in the exit of apical proteins from the Golgi (Iwanami et al., 2016) and Rab11, Rip11, Myosin V and the exocyst complex (e.g. Sec6) are important in the secretion of Rh1 and other rhabdomeral proteins (Satoh et al., 2005; Li et al., 2007; Beronja et al., 2005). Secretory vesicle carrying Rh1 and other rabdomere-destined proteins are moved along actin fibers of the rabdomere terminal web driven by the Myosin V motor (Li et al., 2007). Following the path of Rh1, once at the rhabdomeres, Rab5 and Shibire/Dynamin (Dyn) are required for its endocytosis (Satoh et al., 2005; Alloway et al., 2000; Kiselev et al., 2000). Some of this endocytosed Rh1 is recycled back to the rhabdomere through a retromer-dependent pathway that involves the retromer protein Vps26, whereas the rest is sent to the lysosome for degradation (Wang et al., 2014; Chinchore et al., 2009).

**Figure 1.**
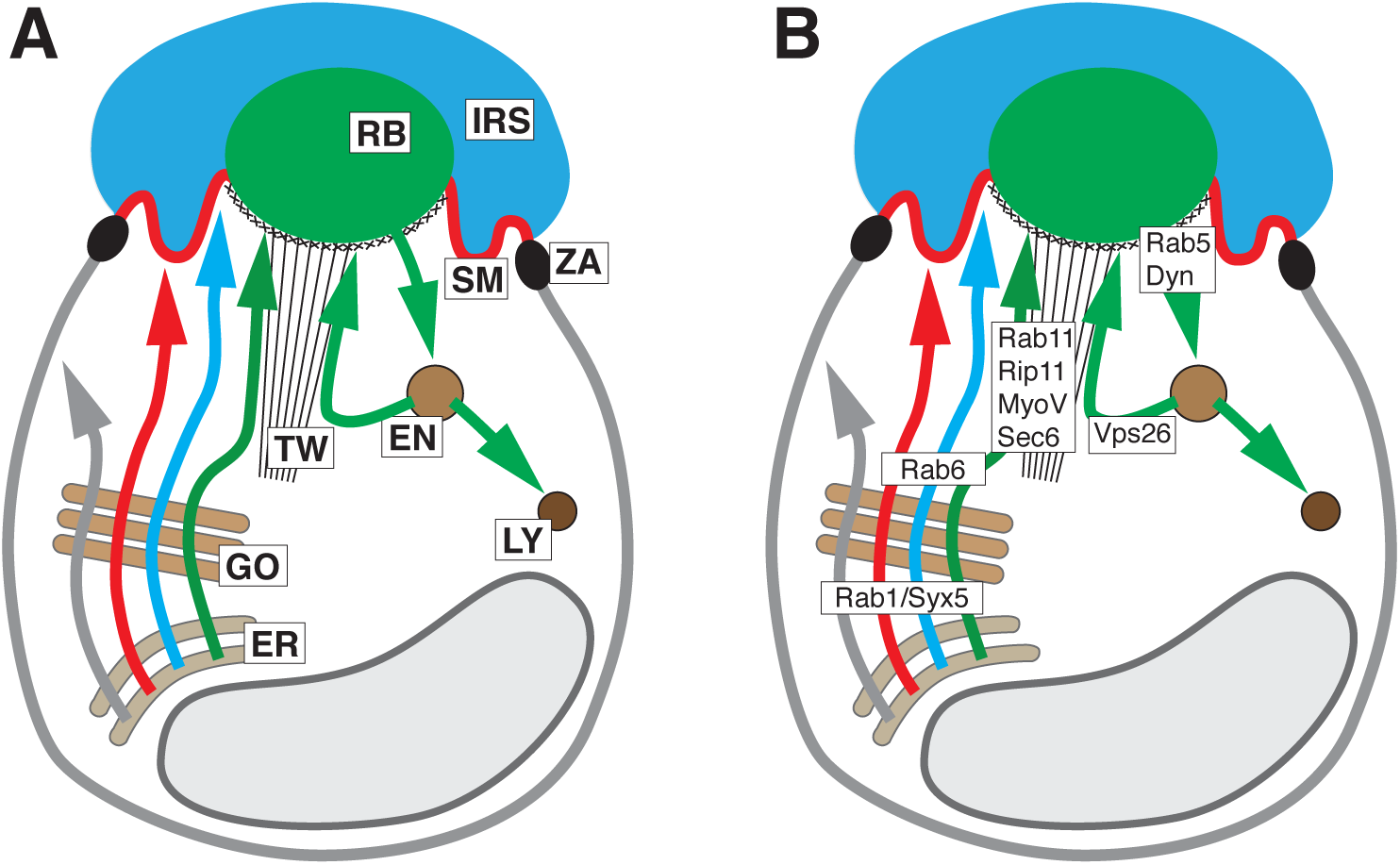
Structure and trafficking of Drosophila PRCs. **(A)** Schematic of PRC showing major known trafficking pathways. Arrows highlight basolateral (grey) and apical (red, Crb; blue, Eys; green, Rh1) trafficking pathways. Abbreviations: EN, endosome; ER, endoplasmatic reticulum; GO, Golgi apparatus; IRS, interrhabdomeral space; LY, lysosome; RB, rhabdomere; SM, stalk membrane; TW, terminal web; ZA, zonula adherens. **(B)** Schematic of PRC showing site of action of major known vesicle trafficking factors. See text for description.

It appears that the apical and basolateral trafficking routes diverge somewhere along the Golgi prior to the action of Rab6, whereas the rhabdomeral versus stalk membrane route diverges downstream of Rab6 following the exit from the Golgi. Crb and Eys are thought to be targeted to the stalk through a pathway distinct from the secretory pathway used by rhabdomeral proteins (Beronja et al., 2005, Husain et al., 2006). The components involved in Crb and Eys exocytosis are currently unknown, except for evidence suggesting that microtubules and microtubule motor proteins are involved in Crb localization during pupal stages (Mukhopadhyay et al., 2010; Chen et al., 2010; League and Nam, 2011; Mui et al., 2011; Nam, 2016).

To develop a better understanding of the apical trafficking mechanisms that control the distribution of Eys, Crb, and Rh1 in the fly retina we analyzed 239 candidate genes. We identified 28 genes that are important for the localization or concentration of apical proteins, and provide a more detail analysis of four factors: The Arf guanine nucleotide exchange factor (GEF) Sec71, the exocyst complex, the microtubule motor dynein, and the endocytotic regulator Syntaxin 7 (Syx7)/Avalanche (Avl).

## Results and Discussion

### Identification of genes involved in PRC vesicle trafficking

Drosophila PRCs are organized as elongated cylinders surrounding a lumen, the IRS, bound by PRC apical membranes (Figure 2A). PRCs deficient in Rh1, Eys, or Crb have prominent developmental defects. Rh1 is essential for rhabdomeral maintenance (Kumar & Ready, 1995) and disruption of factors, such as *Sec6* that affects the exocytosis of Rh1 and other rhabdomeral proteins show rhabdomeral deterioration (Beronja et al., 2005) (Figure 2B). The rhabdomeres of *crb* deficient PRCs appear rectangular in cross-sections as a consequence of a distal to proximal extension defect (Pellikka et al., 2002) (Figure 2B), and *eys* mutant ommatidia lack an IRS (Husain et al., 2006; Zelhof et al., 2006) (Figure 2B).

**Figure 2.**
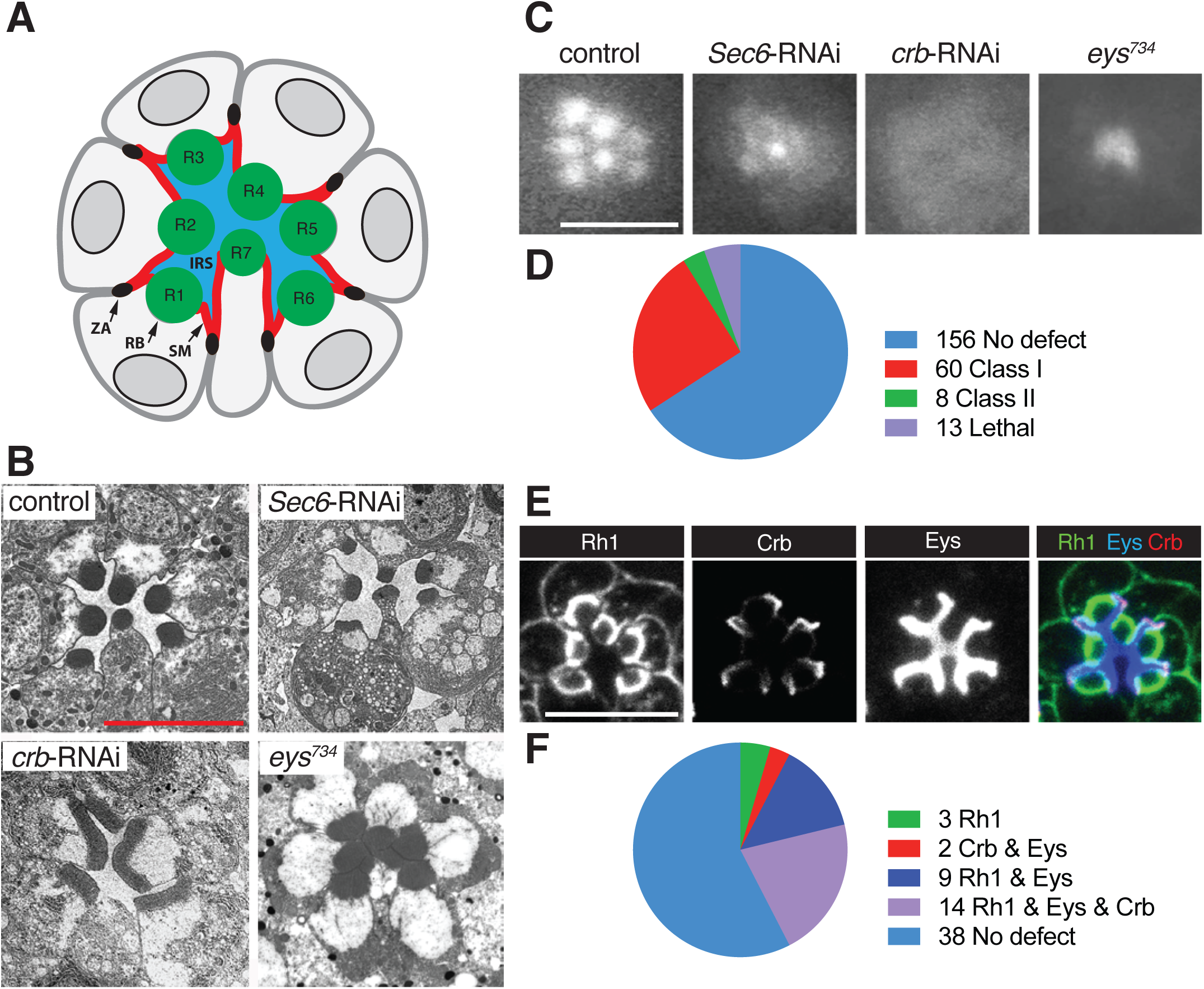
RNAi-based screen of known and predicted trafficking factors in Drosophila PRCs. GMR-GAL4 was used to drive the expression of UAS-dicer-2 and UAS-RNAi constructs. UAS-dicer-2/+; pGMR-Gal4/+ was used as control. Scale bars, 5 µm. **(A)** Schematic of cross-section of PRCs in a Drosophila ommatidium. Apical membranes of PRCs are subdivided into the rhabdomere (RB) and stalk membrane (SM). The stalk membrane connects the rhabdomere to the zonula adherens (ZA). PRCs R1-R8 (R8 is found below R7) surround the interrhabdomeral space (IRS). **(B)** TEM images of a wild-type, *Sec6*-RNAi, *crb*-RNAi, and *eys^734^* mutant ommatidium. **(C)** TLI images of control, *Sec6*-RNAi, *crb*-RNAi, and *eys^734^* mutant ommatidium. **(D)** Summary of the TLI screen. **(E)** Distribution of Rh1, Crb, and Eys in ommatidia of control flies. **(F)** Summary of data obtained from Rh1, CRb, and Eys retinal immunostaining of 68 Class I and Class II candidates identified with TLI (see also Figures S1-S4). Knockdown of 3 genes changed the distribution of Rh1, knockdown of 2 genes changed the distribution of both Crb and Eys, knockdown of 9 genes changed the distribution of both Rh1 and Eys, and knockdown of 14 genes changed the distribution of Rh1, Crb, and Eys.

To identify genes associated with apical trafficking we screened RNAi lines targeting genes known or predicted to be involved in vesicle trafficking for defects similar to Rh1, Crb, and Eys reduction. The eye-specific driver pGMR-Gal4 (Freeman, 1996) was used to drive expression of UAS-dicer (to amplify the effects of RNAi; Ketting et al., 2001) and a total of 291 RNAi lines corresponding to 239 genes (Table S1).

The first step in this screen was conducted using transmitted light illumination (TLI). TLI takes advantage of the fact that rhabdomeres act like optical fiber cables. When a beam of light is transmitted through the eye, individual rhabdomeres become visible, after the cornea has been optically neutralized in an appropriate medium (Franceschini and Kirschfeld, 1971a). Using TLI, 7 distinct rhabdomeres were visible in control flies (Figure 2C). Deviations from the control were categorized into 3 classes based on the severity and type of the defect. Mild to moderate rhabdomeral defects were categorized as class I. Here, one or more, but not all 7 rhabdomeres were distinguishable as individual entities. For example, the defect caused by *sec6* knockdown, which compromises Rh1 trafficking (Beronja et al., 2005), would be categorized as class I (Figure 2C). Ommatidia where rhabdomeres appeared as a single diffuse patch were categorized as class II, as seen for example with *crb* knockdown (Figure 2C). Rhabdomeres do exist in Crb deficient PRCs, but they transmit light poorly, likely as a result of their extension defect (Pellikka et al., 2002). Finally, a class III defect was qualitatively different and referred to an *eys*-like defect, where a single bright spot of light was visible per ommatidium (Figure 2C), as a result of a loss of the IRS (Husain et al., 2006; Zelhof et al., 2006).

Figure 2D summarizes the results of the TLI screen. Out of the 239 genes screened, we found that the knockdown of 69 genes produced a class I or II defect. We did not observe a class III defect. This suggested that from the list of the trafficking proteins tested, no factor exclusively affects Eys trafficking without affecting other apical proteins. Additionally, 15 genotypes were associated with lethality, indicating a leaky expression of the RNAi construct in essential tissues.

RNAi knockdown causing a class I or II defect were further analyzed through immunostaining of retinas for Rh1, Crb, and Eys (Figure 2E). Two phenotypes that were not further analyzed were caused by *Rab11* and *Rab21* knockdown. Knockdown of these genes led to fragile eye tissue that fell apart during dissection. Out of the 67 genes examined, we found that the knockdown of 28 changed the amount and/or localization of one or more apical proteins (Figure 2F and Table 1). We did not find genes that have an exclusive effect on Eys or Crb. We also did not find a case where Crb and Rh1 trafficking was affected in the absence of changes to Eys.

**Table 1.** Summary of RNAi screen. Trafficking genes affecting Rh1, Crb, and Eys distribution in PRCs.

| Cargo | Gene | CG # | RNAi strain | Phenotype |
| --- | --- | --- | --- | --- |
| Rh1 | *Bet3* | CG3911 | BDSC: 38302 | Cytoplasmic Rh1 |
|  | *CdGAPr* | CG10538 | BDSC: 6437 | Cytoplasmic Rh1 |
|  | *twf* | CG3172 | BDSC: 35365 | Cytoplasmic Rh1 |
| Crb & Eys | *sqh* | CG3595 | BDSC: 33892 & 32439 | Larger IRS, longer Crb stained membrane, rectangular rhabdomeres for some ommatidia |
|  | *Rph* | CG11556 | BDSC: 25950 | Larger IRS, longer Crb stained membrane, rectangular rhabdomeres |
| Eys & Rh1 | *Dhc64c* | CG7507 | BDSC: 36698, 28749, 36583 | Increase in cytoplasmic Rh1, cytoplasmic Eys and decrease in IRS size |
|  | *Khc* | CG7765 | BDSC: 25898, 35409 | Cytoplasmic Rh1 and Eys |
|  | *Rab2* | CG3269 | BDSC: 28701 | Missing rhabdomeres and reduced IRS size |
|  | **Exocyst** |  |  |  |
|  | *Exo70* | CG7127 | BDSC: 55234 | Increase in cytoplasmic Rh1, cytoplasmic Eys and decrease in IRS size |
|  | *Exo84* | CG6095 | BDSC: 28712 |  |
|  | *Sec10* | CG6159 | BDSC: 27483 |  |
|  | *Sec15* | CG7034 | BDSC: 27499 |  |
|  | *Sec5* | CG8843 | BDSC: 27526 |  |
|  | *Sec6* | CG5341 | VDRC:105836, 22079 |  |
| Eys & Rh1 & Crb | *Chc* | CG9012 | BDSC: 27530 | Reduced Crb, cytoplasmic Eys and Rh1 |
|  | *aux* | CG1107 | BDSC: 35310 | Reduced Crb, cytoplasmic Eys and Rh1 |
|  | *shi* | CG18102 | BDSC: 36921, 28513 | Cytoplasmic Eys, Crb, and Rh1 and retinal degeneration |
|  | *TSG101* | CG9712 | BDSC: 35710 | Cytoplasmic Rh1, smaller/missing rhabdomeres, smaller IRS and less Crb |
|  | *Slh* | CG3539 | BDSC: 34335 | Reduction of Rh1, Eys and Crb |
|  | *Shrb* | CG8055 | BDSC: 38305 | Cytoplasmic Eys, Crb, and Rh1 and retinal degeneration |
|  | *Vha26*<br>*Vha100-1* | CG1088<br>CG1709 | BDSC: 38996<br>BDSC: 26290 | Cytoplasmic accumulation of Rh1, Crb and Eys |
|  | *Klc* | CG5433 | BDSC: 36795 | Cytoplasmic Rh1 and Eys and reduction in Crb levels |
|  | *Rab1* | CG3320 | BDSC: 27299 | Reduction in Crb, Eys and Rh1 |
|  | *Rab5* | CG3664 | BDSC: 30518 | Many ommatidia disintegrated; less affected ommatidia had larger IRS, longer Crb stained stalks and degenerated rhabdomeres; reduced Rh1 at the rhabdomeres |
|  | *MyosinV* | CG2146 | VDRC: 16902 BDSC: 55740 | Ectopic rhabdomeres at the basolateral membrane. Basolateral Eys and Crb. Cytoplasmic Rh1 and Eys. |
|  | *Avl/Syx7* | CG5081 | BDSC: 29546, VDRC:107264 | Larger IRS, basolateral Rh1, longer Crb stained stalks |
|  | *Sec71* | CG7578 | BDSC: 32366, VDRC:100300 | Significant reduction in Crb, Eys and Rh1, rhabdomeres absent |

We performed a more in-depth analysis of the defects caused by loss of Sec71 (Figure 3), exocyst (Figure 4), dynein (Figure 5) and Syx7/Avl (Figure 6). We selected these factors as their knockdown was associated with robust defects, and as they represented a component of the Golgi (Sec71), secretory (exocyst and dynein), and the endocytic (Syx7/Avl) pathways. Defects caused by the knockdown of other factors can be found in Figure S1 (for RNAi constructs affecting Rh1), Figure S2 (for RNAi constructs affecting Rh1 and Eys), Figure S3 (for RNAi constructs affecting Crb and Eys), and Figure S4 (for RNAi constructs affecting all three apical proteins).

**Figure 3.**
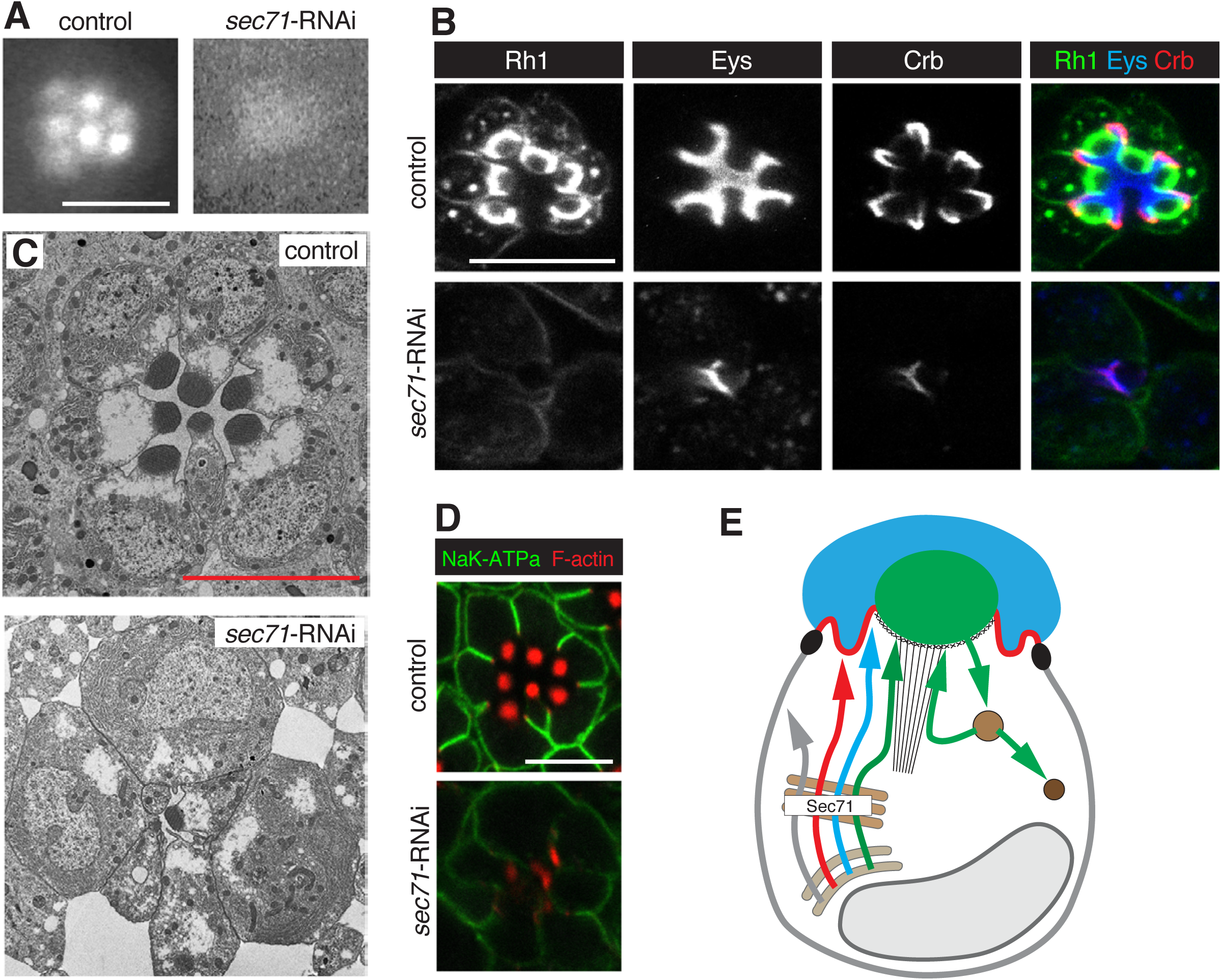
Knockdown of *Sec71* caused a reduction of both apical and basolateral proteins. RNAi line BDSC 32366 (*Sec71*-RNAi) was used. Flies with *Sec71*-RNAi were crossed to UAS-dicer-2; pGMR-Gal4. UAS-dicer-2/+; pGMR-Gal4/+ was used as control. Scale bars are 5 µm. **(A)** TLI does not reveal individual rhabdomeres in *Sec71* knockdown PRCs (Class II defect). **(B)** *Sec71* deficient PRCs show a severe reduction of Rh1, Crb, and Eys. **(C)** TEM shows a loss of IRS, missing rhabdomeres and gaps between ommatidial units of Sec71 deficient PRCs. **(D)** The K^+^Na^+^ATPase subunit Nrv is reduced in *Sec71* knockdown PRCs. Acti-stain555 (F-actin) was used to visualize the rhabdomeres. **(E)** Summary model suggesting that Sec71 acts in the Golgi to control secretion of apical and basolateral factors. See Figure 1A for annotation and text for discussion.

**Figure 4.**
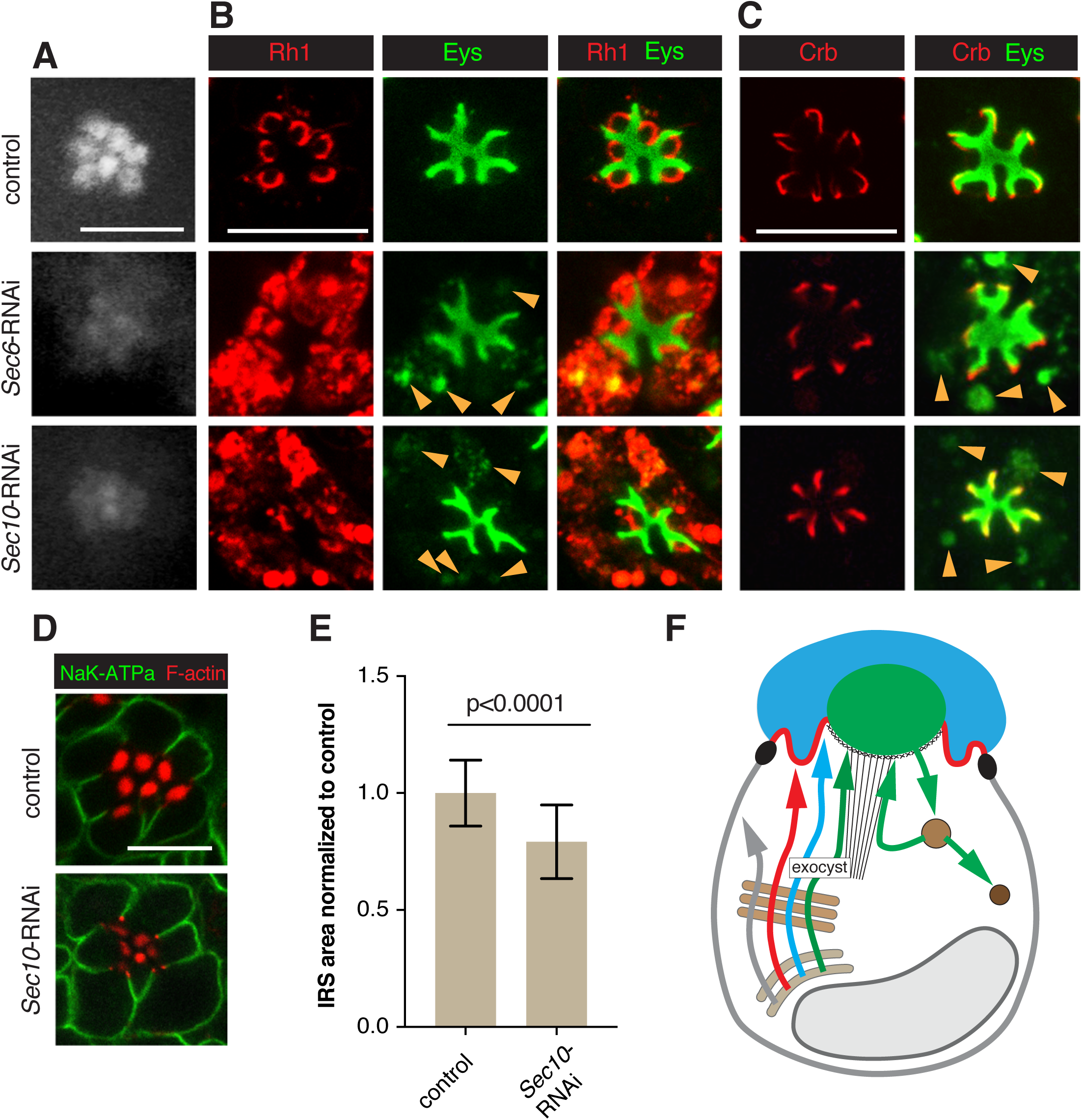
The exocyst contributes to Rh1 and Eys exocytosis. RNAi line VDRC 22079 (*Sec6*-RNAi) was used to knock down Sec6, and line BDSC 27483 (*Sec10*-RNAi) to deplete Sec10. *Sec6*-RNAi or *Sec10*-RNAi were crossed with UAS-dicer-2; pGMR-Gal4. UAS-dicer-2/+; pGMR-Gal4/+ was used as control. Scale bars, 5 µm. **(A)** Individual rhabdomeres were only partially visible in Sec6 and Sec10 deficient retinas using TLI. Both were categorized as Class I. **(B)** Sec6 and Sec10 deficient PRCs show cytoplasmic accumulation of Rh1 and Eys (arrowsheads). Cytoplasmic Eys colocalizes with Rh1. **(C)** No defect was observed for Crb levels or localization in exocyst knockdown PRCs. Cytoplasmic Eys in exocyst deficient PRCs is visible in the absence of Rh1 staining (arrowheads), eliminating the possibility of cross-reaction between Eys and Rh1 antibodies as an explanation for the cytoplasmic Eys signal seen in (B). **(D)** Levels/localization of the basolateral protein Nrv (K^+^Na^+^ATPase subunit) was not affected in exocyst compromised PRCs. Acti-stain555 (F-actin) was used to visualize the rhabdomeres. **(E)** IRS size was significantly reduced in *Sec10* deficient retinas. A total of 69 individual IRS were measured for control and 118 for *Sec10* RNAi ommatidia, using three different animals per genotype. Values were normalized to the control. Error bars represent standard deviation. Unpaired non-parametric Mann-Whitney test. **(E)** Summary model indicating the secretion of Rh1 and Eys containing secretory vesicles depends on the exocyst. See Figure 1A for annotation and text for discussion.

**Figure 5.**
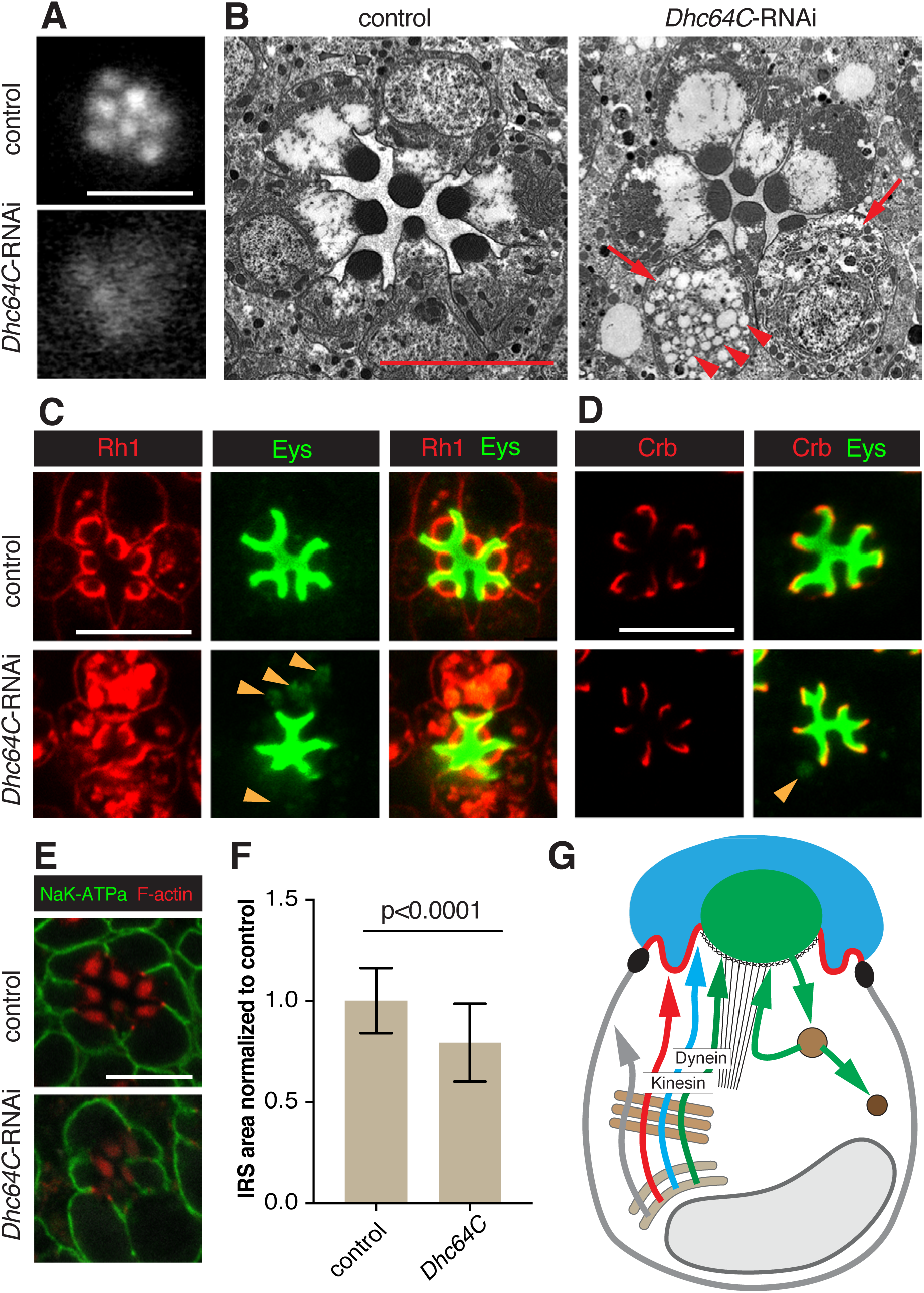
Dynein is important for Rh1 and Eys trafficking. RNAi line BDSC 36583 (*Dhc64C*-RNAi) was used to deplete Dhc64c. *Dhc64C*-RNAi was crossed with UAS-dicer-2; pGMR-Gal4. UAS-dicer-2/+; pGMR-Gal4/+ was used as control. Scale bars, 5 µm. **(A)** Individual rhabdomeres were partially visible in dynein (Dhc64c) deficient retinas. We categorized this defect as Class I. **(B)** TEM analysis revealed a smaller IRS, smaller rhabdomeres, bloated PRCs (arrow) and cytoplasmic accumulation of vesicles (arrowheads) in dynein deficient retinas. **(C)** Dynein deficient PRCs showed cytoplasmic accumulation of Rh1 and Eys (arrowheads). Cytoplasmic Eys colocalized with cytoplasmic Rh1. **(D)** Levels or localization of Crb was normal in Dhc64c depleted PRCs. Arrowhead points to cytoplasmic accumulation of Eys. **(E)** Levels/localization of the basolateral protein Nrv (K^+^Na^+^ATPase subunit) was not affected by dynein depletion. Acti-stain555 (F-actin) was used to visualize the rhabdomeres. **(F)** IRS size was significantly reduced in dynein compromised retinas. A total of 106 individual IRS were measured for control and 109 for dynein deficient ommatidia using 3 different animals per genotype. Values were normalized to the control. Error bars represent standard deviation. Unpaired non-parametric Mann-Whitney test. **(F)** Summary model indicating the secretion of Rh1 and Eys containing secretory vesicles depends on dynein and that secretion of Rh1, Eys, and Crb requires kinesin function (see Figures S2 and S4). See Figure 1A for annotation and text for discussion.

**Figure 6.**
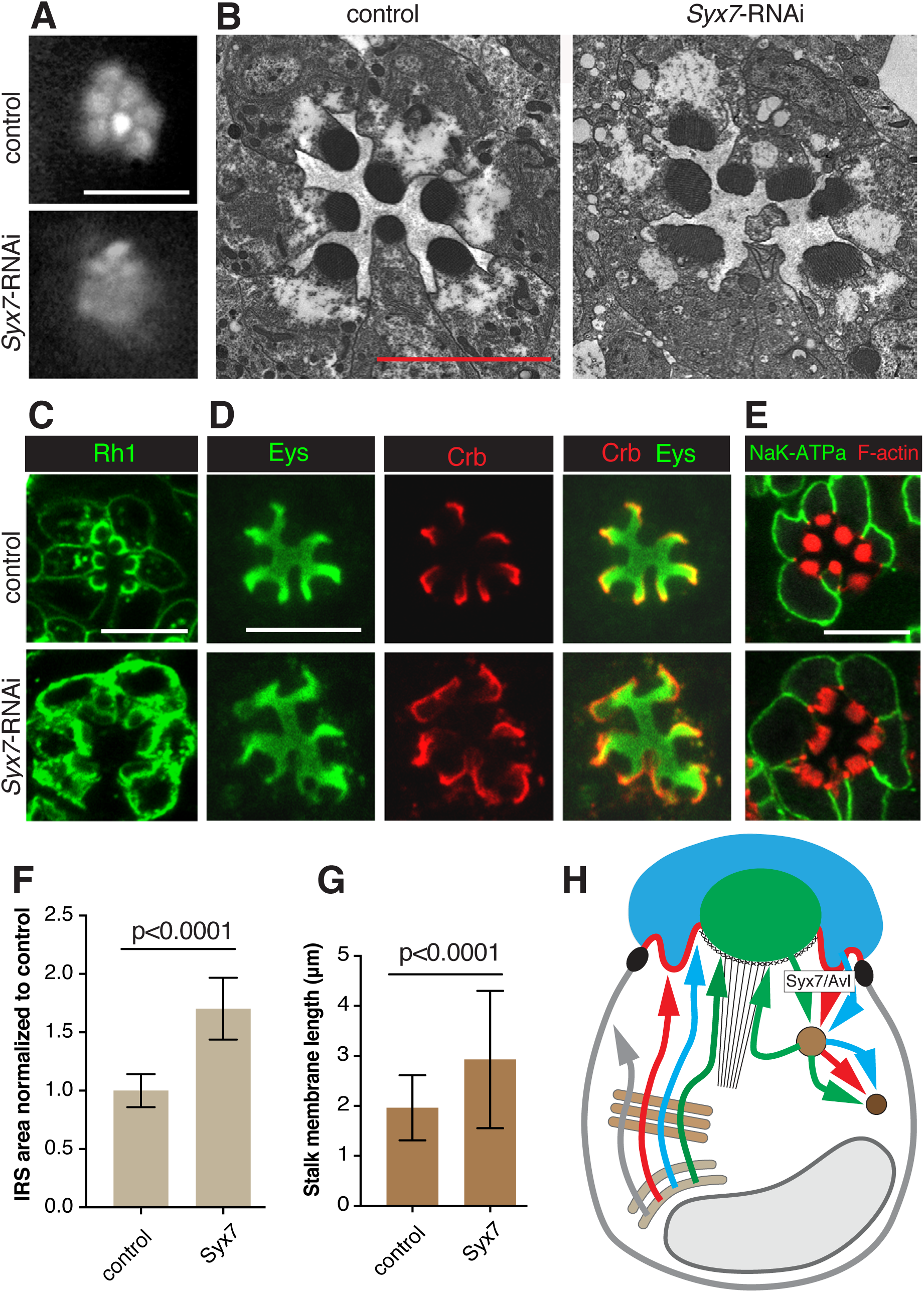
Syx7/Avl deficient PRCs show enhanced surface levels of Rh1, Eys and Crb. RNAi line BDSC 29546 (*Syx7*-RNAi) was used to deplete Syx7/Avl. *Syx7*-RNAi was crossed with UAS-dicer-2; pGMR-Gal4. UAS-dicer-2/+; pGMR-Gal4/+ was used as control. Scale bars, 5 µm. **(A)** Individual rhabdomeres were partially visible in Syx7/Avl deficient retinas. We categorized this defect as Class I. **(B)** Knockdown of *Syx7/Avl* led to rhabdomeral fragmentation and degeneration. **(C)** Syx7/Avl deficient PRCs show increased apical and basolateral accumulation of Rh1. **(D)** IRS area stained with Eys and stalk membranes labeled with Crb are larger in Syx7/Avl deficient PRCs compared to the control. **(E)** Levels/localization of the basolateral protein Nrv (K^+^Na^+^ATPase subunit) is normal in Syx7/Avl deficient PRCs. Acti-stain555 (F-actin) was used to visualize rhabdomeres which appear disorganized. **(F)** IRS size was significantly larger in Syx7/Avl deficient retinas compared to controls. A total of 69 individual IRS were measured for the control and 138 for *Syx7/Avl* knockdown PRCs using 3 different animals per genotype. Values were normalized to the control. Error bars represent standard deviation. Unpaired non-parametric Mann-Whitney test. **(G)** Stalk membranes were significantly larger in Syx7Avl deficient PRCs compared to controls. A total of 182 individual stalk membranes were measured for the control and 126 for Syx7/Avl deficientknockdown PRCs. Error bars represent standard deviation. Unpaired non-parametric Mann-Whitney test. **(H)** Summary model indicating that the endocytosis of Rh1, Crb, and Eys depends on Syx7/Avl. See Figure 1A for annotation and text for discussion.

### Sec71 is required for biosynthetic traffic of both apical and basolateral proteins

The Arf GEF Sec71 is essential for the integrity of Golgi compartments in Drosophila ddaC sensory neurons (Wang et al., 2017). The function of Sec71 in Drosophila PRCs was unknown. The expression of two different *Sec71* RNA lines led to a class II defect as seen with TLI (Figure 3A). We observed a robust reduction but normal distribution of Eys, Crb, and Rh1 (Figure 3B). *Sec71* knockdown affected the integrity and survival of cells in the retina as ultrastructural analysis showed retinal holes, dying and/or absent PRCs, and small or missing rhabdomeres (Figure 3C).

A similar reduction in Eys, Crb, and Rh1 amounts were also observed in Rab1 deficient PRCs (Figure S4K). Rab1 has been associated with the ER to Golgi trafficking of Rh1 (Satoh et al., 1997). Proper processing done at the Golgi is thought to be important for protein stability, and disruption of factors required for ER to Golgi processing such as Syn5 and Rab1 are known to cause an overall reduction in cargo levels (Satoh et al., 2016a, Satoh et al., 1997). Therefore, the reduction of Eys, Crb, and Rh1 seen in Sec71 compromised PRCs may be due to a lack of Golgi processing, suggesting a role for Sec71 at or prior to the Golgi in the biosynthetic pathway.

This conclusion is further supported by our observations that also the basolateral protein Nervana (Nrv), a subunit of the Na^+^/K^+^ ATPase was reduced in *Sec71* knockdown PRCs (Figure 3D). This suggests that Sec71 acts prior to the separation of apical and basolateral trafficking routes, which is thought to occur at or prior to the trans-Golgi network (TGN). Similar defects in apical and basolateral protein transport have been described for the Golgi-associated SNARE protein Syn5 (Satoh et al. 2016a). Taken together, our observations indicate that Sec71 contributes to the secretion of both apical and basolateral proteins in PRCs suggesting a defect in Golgi processing (Figure 3G).

Sec71 is related to the human Arf GEF BIG1/BIG2 family (Christis and Munro, 2012). BIG1 and BIG2 are associated with the TGN (Mansour et al., 1999; Yamaji et al., 2000; Shinotsuka et al., 2002; Zhao et al., 2002; Ishizaki et al., 2008), and the recycling endosomes (Shin et al., 2004; Shen et al., 2006; Ishizaki et al., 2008). In Drosophila sensory neurons, Sec71 is mainly found at the TGN, and the Golgi is severely disrupted in Sec71-deficient cells (Wang et al., 2017).

As an Arf GEF, Sec71 is likely to activate one or more resident Golgi Arfs in the Drosophila PRCs. Candidates to consider are Arf1, Arf4, and Arl1. Arf1 (Arf79F in Drosophila), is important in regulating traffic at the Golgi apparatus (Rodrigues and Harris, 2017). In Drosophila sensory neurons, Sec71 activates Arf1, and the Golgi apparatus is disrupted in *Arf1* mutant cells (Wang et al., 2017). Arf4 (Arf102F in Drosophila), is associated with the TGN and is important in targeting rhodopsin to the cilia of frog PRCs (Mazelova et al., 2009; Wang et al., 2012). Finally, Arl1 (Arf72A in Drosophila), localizes to the Golgi in Drosophila PRCs and is important in quality check of cargo (Lee et al., 2011). Loss of Arl1 results in an increase of Golgi compartments and an acceleration of Rh1 secretion (Lee et al., 2011). In our TLI screen, the expression of Arf1 and Arl1 RNA lines did not show an effect. This may be due to the inactivity/low activity of the RNAi lines used, or functional redundancies between PRC Arfs. The expression of Arf4 dsRNA led to lethality, which did not allow us to further examine its function in the eye.

### The exocyst complex contributes to both Rh1 and Eys secretion

Depletion of exocyst components Exo70, Exo84, Sec10, Sec15, Sec5 and Sec6 led to class I defects (see Figure 4A for Sec6 and Sec10 data). Rhabdomeral defects were expected as the exocyst is known to be important for the exocytosis of rhabdomeral proteins (Beronja et al., 2005). We observed cytoplasmic accumulation of Rh1 in *Sec6* and *Sec10* knockdown PRCs as previously described for *Sec6* mutant PRCs (Beronja et al., 2005) (Figure 4B), smaller rhabdomeres and a cytosolic accumulation of secretory vesicles (Figure 2B).

Unexpectedly, in addition to Rh1 trafficking defects, we also observed cytoplasmic Eys for all exocyst RNAi lines tested (see Figure 4B and C). Cytoplasmic Eys and Rh1 partially overlapped with all cytoplasmic Eys accumulations also highly enriched in Rh1, suggesting that Eys and Rh1 are trapped in the same cytoplasmic compartments in exocyst deficient PRCs (Figure 4B). Expression of several exocyst RNAi constructs also resulted in a smaller than normal IRS (Figure 4B). We quantified the size of the Eys-positive area in Sec10 depleted PRCs in comparison to control ommatidia and found that the IRS size was reduced by ~20% (Figure 4E). As IRS size is directly dependent on Eys level (Husain et al., 2006; Zelhof et al., 2006), it is likely that the IRS is reduced due to ineffective Eys trafficking. We did not observe a noticeable change in Crb (Figure 4C) or basolateral trafficking of Nrv (Figure 4D).

It was previously thought that Eys was secreted through the stalk membrane and would not be affected by defects in rhabdomeral trafficking (Husain et al., 2006). The different impact on Eys trafficking observed here compared to our previous analysis may be the result of the different strategies used to compromise exocyst function (RNAi used here versus generation of *Sec6* mutant clones combined with the expression of a partial rescue construct to avoid cell lethality caused by the loss of exocyst function; Beronja et al., 2005; Husain et al., 2006). We conclude that the exocyst complex, which is crucial for trafficking to the rhabdomere, is also important for Eys trafficking to the IRS (Figure 4F).

### Microtubule motor protein dynein is required for Rh1 and Eys secretion

The function of dynein, a minus-end directed microtubule motor protein, has not been previously described in fly PRCs. We observed class I defects with three distinct RNAi lines targeting Dynein heavy chain 64C (Dhc64C) (Figure 5A). We observed abnormal cytoplasmic accumulation of Rh1 and Eys for all three RNAi lines (Figure 5C,D). The presence of cytoplasmic Eys partially overlapped with Rh1, suggesting that Eys and Rh1 are trapped in the same compartments. Cytoplasmic Eys was coupled with a reduction in IRS size (Figure 5F). We did not observe an effect on Crb (Figure 5D) or basolateral Nrv (Figure 5E), suggesting that traffic to the stalk and the basolateral membrane was not affected.

Expanded cell bodies and an accumulation of cytoplasmic vesicles of various sizes are apparent in Dhc64C depleted PRCs (Figure 5B). These vesicles looked similar to the secretory vesicles described for PRCs compromised for the exocyst, Rab11, Rip11, or Myosin V (Satoh et al., 2005; Beronja et al., 2005; Li et al., 2007). Noteably, we also observed similar defects with the microtubule plus-end directed kinesin motor. Depletion of Kinesin heavy chain (Khc) and Kinesin light chain (Klc) led to a Class I defect in TLI. Rh1 and Eys accumulated in the cytoplasm and IRS size was decreased and depletion of Klc but not Khc led to a reduction in Crb at the stalk membrane (Figure S2 and S4). It is curious that the knockdown of both plus-end and minus-end directed motors caused similar Eys and Rh1 secretion defects.

Klc deficient PRCs also showed a reduction in Crb (Figure S1J). Microtubules and microtubule-related proteins are known to be important for Crb trafficking in the pupal PRCs (Mukhopadhyay et al., 2010; Chen et al., 2010; League and Nam, 2011; Mui et al., 2011; Nam, 2016). It is not clear whether this Crb defect is specific to Klc knockdown or if a stronger knockdown of Khc or Dhc64C would affect Crb in a similar manner.

Previous studies in the developing eye discs had shown that the plus-end of microtubules are found at the apical cell membrane whereas the minus-ends are located in the vicinity of the cell nucleus (Mosley-Bishop et al. 1999). Microtubule orientation in adult PRCs is not known and would be important to address in future studies to help in the interpretation of defects resulting from knockdown of motor proteins. It is known that vertebrate PRCs possess microtubules with opposing orientations (for review see Nemet et al., 2015). In vertebrate PRCs, rhodopsin is made in the inner segment (IS) and transported to the vertebrate equivalent of the rhabdomere, the outer segment (OS). Within the IS, microtubules are orientated with their plus-ends at the Golgi and minus-ends at the base of the OS. Microtubule orientation is reversed in the OS, where the minus-end is found at the base of the OS, while the plus-end is found at the distal end of the OS. It has been proposed that dynein is important in the trafficking of rhodopsin from Golgi to the OS, whereas kinesin is important in trafficking from the base of the OS to its tip (Nemet et al., 2015). Loss of dynein or kinesin function may therefore lead to defects in rhodopsin trafficking.

Taken together, we found that microtubule motors, kinesin and dynein are essential in the secretion of Eys and Rh1. One important conclusion from our findings is that microtubule and actin-based transport mechanisms cooperate in the delivery of rhabdomere-targeted proteins. Previous studies had revealed the importance of actin filament-based routes and the actin motor protein Myosin V in rhabdomeral vesicle trafficking (Li., et al., 2007). It would be interesting to examine how microtubule and actin-based transport mechanisms interface to facilitate apical trafficking in PRCs.

### Syntaxin 7/Avalanche is required for Rh1, Eys, and Crb endocytosis

Syx7/Avl is required for apical endocytosis in imaginal disc epithelia. Syx7/Avl deficient cells displayed an excessive accumulation of Crb at the apical membrane (Lu and Bilder, 2005). Syx7/Avl co-localizes with Rab5 and is required for the entry of cargo proteins, such as Crb into early endosomes (Lu and Bilder, 2005). Previous results suggested that endocytosis is essential for rhabdomeral maintenance in Drosophila PRCs. Endocytosis is thought to be important in the proper formation of the interface between the rhabdomere and the sub-rhabdomeric space where the rhabdomere terminal web in located (Pinal and Pichaud, 2011). Here, we found that knockdown of Syx7/Avl using two distinct RNAi lines led to a class I defect with TLI (Figure 6A), and consistent with earlier results indicating that endocytosis promotes rhabdomere integrity, we observed structural defects in rhabdomeres of Syx7/Avl knockdown PRCs (Figure B,E).

Rh1, Crb, and Eys levels were increased in Syx7/Avl compromised PRCs whereas Nrv levels remained normal (Figure 6C,D,E). Corresponding to the increase in Eys (Zelhof et al., 2006) we found an enlarged IRS in Syx7/Avl knockdown PRCs (Figure 6F). Similarly, the overabundance of Crb in Syx7/Avl knockdown PRCs was associated with an enlarged stalk membrane (Figure 6G) as previously reported for PRCs that overexpress Crb (Pellikka et al., 2002). The apical accumulation of Eys and Crb as a result of compromised apical endocytosis suggests that both proteins similar as Rh1 undergo active turn-over in Drosophila PRCs.

Interestingly, Rh1 was increased not only at the apical rhabdomere but also at the basolateral membrane in Syx7/Avl knockdown PRCs, where in control PRCs only small amounts of Rh1 are found (Figure 6C). It is possible that an over-accumulation of Rh1 at the rhabdomeres led to a leakage of Rh1 into the basolateral membrane. Alternatively, it is possible that Rh1 may normally be transcytosed from the basolateral to the apical membrane, requiring basolateral endocytosis. As we did not find an effect on the basolateral protein Nrv (Figure 6E) we favor the first possibility, and would predict that Syx7/Avl is associated with the apical membrane of PRCs.

### Concluding remarks

By exploring apical vesicle trafficking in the fly PRCs through an RNAi-based screen we have uncovered 28 genes involved in apical localization of the key PRC proteins Rh1, Crb, and Eys. We have shown that the Arf GEF Sec71 is essential for proper apical and basolateral protein trafficking, the exocyst complex and microtubule motors dynein and kinesin are important for Eys and Rh1 secretion, and the syntaxin Avl is involved in Crb, Eys, and Rh1 endocytosis. Our results have implications for the understanding of human eye diseases as mutations in human orthologs of Eys, Crb, and Rh1 cause eye degenerative diseases in humans (Abd El-Aziz et al., 2008; Collin et al., 2008; Richard et al., 2006; den Hollander et al., 2008; Hollingsworth and Gross 2012). The rhabdomeres and the stalk membrane in Drosophila correspond topologically and functionally to the vertebrate rod and cone outer segment and inner segment, respectively. It is likely that similar factors are involved in apical trafficking in Drosophila and vertebrate PRCs. One example is Rab11, which was shown to be important in post Golgi Rh1 exocytosis in both Drosophila and vertebrates (Satoh et al., 2005; Mazelova et al., 2009). It will therefore be of interest to further explore the conservation of vesicle trafficking mechanisms in Drosophila and mammalian PRCs.

## Materials and Methods

### Fly stocks and crosses

UAS-RNAi lines were obtained from the Bloomington Drosophila Stock Center (BDSC) and Vienna Drosophila Resource Center (VDRC). See Table S1 for a list of lines used. UAS-RNAi constructs were expressed in the developing eye with UAS-dicer-2; GMR-Gal4. GMR-Gal4 activity starts in the developing retina posterior to the morphogenetic furrow at third larval instar and continues to adulthood (Freeman, 1996). UAS-dicer-2 was used to amplify the effects of RNAi (Ketting et al., 2001). Flies were raised at 25°C under dark conditions. 0-24 hour old adult progeny were analyzed. We used UAS-dicer-2/+; pGMR-Gal4/+ as control for all experiments.

### Transmitted Light Illumination (TLI)

Heads of 0-24 hour old adult flies were detached and glued in an anterior up orientation to a microscope slide using a thin layer of clear nail polish. A drop of oil (Immersol, 518N) was placed on the sample and the eyes were examined under an upright light microscope (Zeiss Axiophot2). Shining a narrow but bright beam of light from below the sample and optically neutralizing the cornea in an appropriate medium made the rhabdomeres visible (Franceschini and Kirschfeld, 1971a).

### Immunohistochemistry

The retinas of 0-24 hour old adult flies were dissected in phosphate buffer (pH 7.4). The cornea and the rest of the head including the brain tissue were removed using forceps. Subsequently, the retinas were fixed in 4% formaldehyde in phosphate buffer (pH 7.4) for 15 minutes, followed by 30 minutes wash in phosphate buffer (pH 7.4). Prior to antibody staining, the retinas were kept in a 0.3% Triton X-100 phosphate buffer at 4°C for 24 hours or longer. The antibody staining was done according to a standard protocol. The following antibodies were used: rat anti-Crb (F3, 1:500; Pellikka et al., 2002); mouse anti-Rh1 (4C5, 1:50; Developmental Studies Hybridoma Bank), guinea pig anti-Eys (G5, 1:500; Husain et al., 2006), mouse anti-Nervana (nrv5f7, 1:50, Developmental Studies Hybridoma Bank), rabbit anti-GM130 (ab30637, 1:300, abcam). Secondary antibodies anti-guinea pig alexa fluor 647 (A21450), anti-rat alexa fluor 555 (A21434), and anti-mouse alexa fluor 488 (A11029) were used at 1:400 (Molecular Probes/Thermo Fisher Scientific). Acti-stain555 (PHDG1-A, 1:75, Cytoskeleton Inc) was used to visualize the rhabdomeres. A Leica TCS SP8 confocal microscope with 100x objective was used to capture images. Image J (Fiji) and Adobe Photoshop CS5.1 were used to edit and compile figures.

### Electron Microscopy

Transmission electron microscopy (TEM) was performed on 0-24 hour old adult flies. Detached heads were bisected in ice-cold fixative solution (2% para-formaldehyde and 2.5% glutaraldehyde in 0.1 M sodium cacodylate, pH 7.4). The dissected tissue was kept in the above-mentioned fixative for 3 days at 4°C on a nutator. Next, the tissue was washed with 0.1 M sodium cacodylate and treated with a solution of 1% osmium tetroxide and 0.1 M sorbitol in 0.1 M sodium cacodylate for 1 hour in the dark. After this treatment, the eyes were again washed with 0.1 M sodium cacodylate and dehydrated in a series of low to high concentrations of ethanol (50%, 70%, 80%, 100%) and were then infiltrated and imbedded in Spurr’s resin. Ultra-thin sections were stained with uranyl acetate and lead citrate. A Hitachi HT7000 transmission electron microscope was used to view the tissue at 700x magnification and an AMT XR-111 digital camera with AMT capture engine software (version 5.03) was used to capture images.

### Interrhabdomeral space (IRS) quantification

Retinas were immuno-stained with anti-Eys antibody. Images were taken using a Leica TCS SP8 confocal microscope, with a 100x objective (NA 1.4). IRS size was then measured using Imaris software. Mean and standard deviation were calculated. An unpaired non-parametric Mann-Whitney test was performed to establish p values.

### Stalk membrane quantification

Image J (Fiji) was used to measure stalk membranes from TEM images taken at 700x magnification. Individual stalk membranes were traced from the base of the rhabdomeres to the ZA and the length was measured. Mean and standard deviation were calculated. An unpaired non-parametric Mann-Whitney test was performed to establish p values.

### Data and reagent availability

All data are included in the paper or the associated supplemental materials. All Drosophila stocks are available from public repositories. All other reagents are commercially available or can be sent upon request.

## Acknowledgements

We like to thanks Milena Pellikka technical assistance. We are grateful for the support of Audrey Darabie and the Imaging Facility of the department of Cell and Systems Biology. We thank Dorothea Godt for critical reading of the manuscript and helpful suggestions. A.L. was supported by a NSERC (CGS) Alexander Graham Bell fellowship. The work was supported by the Canadian Institutes for Health Research.

## Author contribution

Experimental work: A.L.

Experimental design and evaluation: A.L. and U.T. Fundraising and supervision: U.T.

Preparation of manuscript: A.L. and U.T.

The authors declare no competing interests.

